# Smooth muscle Cxcl12 activation is associated with vascular remodeling in flow-induced pulmonary hypertension

**DOI:** 10.1101/2024.09.10.611870

**Authors:** Timothy Klouda, Savas T. Tsikis, Thomas I. Hirsch, Yunhye Kim, Tiffany Liu, Ingeborg Friehs, John Y.-J. Shyy, Gary Visner, Benjamin A Raby, Mark Puder, Ke Yuan

## Abstract

Patients with congenital heart disease (CHD) resulting in significant left-to-right shunting of blood are at risk for the development of pulmonary arterial hypertension (PAH). The underlying mechanism by which pulmonary overcirculation and shear stress lead to vascular remodeling remains unclear. Our study established a new “two-hit” murine model of severe pulmonary hypertension (PH) by combining left pneumonectomy and exposure to hypoxia (LP/Hx). Utilizing transgenic reporter lines, immunofluorescence staining, and advanced microscopy, we conducted cell-lineage tracing experiments for endothelial cells (ECs), smooth muscle cells (SMCs), and pericytes. We identified that SMCs is a primary contributor to distal arteriolar remodeling after LP/Hx. Subsequent qPCR analysis on isolated cells demonstrated that *Cxcl12* was upregulated in both ECs and SMCs from LP/Hx animals. Likewise, CXCL12 was overexpressed in the SMC layer of arterioles in patients with acyanotic PAH-CHD. These findings provide novel insights into the contribution of SMCs and Cxcl12 to pulmonary flow-induced vascular remodeling. This newly established murine model of PH will serve as a new tool for research and targeted therapeutics for patients with PAH.

## Introduction

Pulmonary arterial hypertension (PAH) is a devastating, chronic, and progressive disease characterized by remodeling of the distal arterioles and elevated pulmonary artery pressures (PAPs)^1,2^. A significant histopathological feature of the vasculature in patients with PAH is neointimal hyperplasia of the distal arterioles, which involves dysregulated proliferation of smooth muscle cells (SMCs) and apoptosis-resistant endothelial cells (ECs)^3^. Pulmonary overcirculation is implicated in vascular remodeling and can lead to the development of pulmonary hypertension (PH). Notably, 37.9% of patients develop a mild/moderate increase in PAPs after major lung resection, and 3.4% develop severe PH^4-6^. Increased pulmonary blood flow and shear stress often result from intra- or extra-cardiac shunting, especially in patients with specific types of congenital heart disease (CHD) leading to pulmonary overcirculation. Approximately 3-10% of patients with CHD, particularly those with large left-to-right shunts such as a ventricle septal defect (VSD) or atrioventricular canal defect are at risk of developing PAH^7-9^. Despite advances in the care of patients with CHD, there are no disease-modifying therapies for PAH-CHD and its diagnosis is associated with increased healthcare costs, morbidity, and mortality^7-12^.

The mechanisms resulting in vascular remodeling associated with pulmonary overcirculation remain unclear^9,13^. Early-stage PAH-CHD is typically reversible and the histological changes are characterized by medial hypertrophy and increased muscularization of the distal arterioles, while late-stage disease leads to permanent changes in the vasculature including neointimal formation and the development of plexiform lesions^14-16^. Studies investigating the pathogenesis of PAH-CHD and the effects of shear stress on vascular remodeling have primarily focused on the response and contribution of ECs^16-18^. The increased hemodynamic stress exerted on the pulmonary arterioles disrupts the endothelium, activating mechanotransductive mechanisms that contribute to the development of PH^19^. A “two-hit” model of PH, combining left pneumonectomy (LP) with the VEGFR antagonist Sugen-5416 (SUGEN) or monocrotaline (MCT) (MCT pyrrole or MCTP for mice), has been employed to study flow-induced PH, predominantly in rats^20-24^. However, these rodent models of PH have limitations, including concurrent compensatory lung growth (CLG) and the inability to recapitulate the physiological conditions experienced by patients. An animal model of PH combining increased blood flow and hypoxia (Hx) may better represent the physiological conditions of patients, enabling a comprehensive study of the underlying pathogenesis and the evaluation of potential disease-modifying therapies.

Chronic Hx is often employed to induce PH in rodents. Murine models of Hx-induced PH are characterized by an increased presence of SMA+ cells on the distal arterioles^25-28^. Similarly, a distinct histological feature of arterioles from patients with PAH is the increased accumulation of SMA+ SMCs in vascular lesions^29^. Patients with CHD can be exposed to prolonged periods of Hx due to sleep disorder breathing (SDB), associated cardiopulmonary anomalies, infections, and during operative procedures^30-32 33-35^. When exposed to Hx, pulmonary blood flow is redirected to better-ventilated regions of the lung by contraction of pulmonary artery SMCs, which can result in increased pulmonary vascular resistance and tone (known as hypoxic pulmonary vasoconstriction)^36,37^. The potential synergistic effects of Hx and shear stress on vascular remodeling, as well as the molecular pathways contributing to flow-induced PH, remain largely under investigation.

CXCL12 is a chemokine that coordinates cellular responses to injury, inflammation, and angiogenesis. Inhibition of the downstream signaling pathway through its receptor, CXCR4, leads to decreased SMC proliferation and pericyte coverage on arterioles^38,39^. We previously demonstrated that Cxcl12 overexpression drove NG2+ pericytes to gain a SMC-like phenotype and participate in vascular remodeling in Hx-induced PH^40,41^. Additionally, the upregulation of Cxcl12 has been documented in multiple animal models of PH, including SUGEN/Hx rats, MCT rats, Hx bovine, Hx mice, and prolyl hydroxylase domain-containing protein 2 (PHD2) or von Hippel–Lindau loss of function mice^42-50^. Furthermore, in PAH clinical samples, ECs, pericytes, and adventitial fibroblasts display augmented CXCL12 activity^41,47,51^. Despite the growing literature and evidence on the role of Cxcl12 in the development of Hx-induced PH, its role in flow-induced vascular remodeling is unknown.

In this study, we introduce an augmented murine model of severe PH by combining increased pulmonary blood flow via LP and chronic Hx. Our key findings are as follows: 1) A new murine model of PH with high right ventricle systolic pressure (RVSP), right ventricle hypertrophy (RVH), and increased vascular remodeling of the distal arterioles; 2) Using a transgenic reporter mouse line (*Acta2-CreER::R26-mTmG)*, SMCs are a key contributor to vascular remodeling after LP and LP/Hx; 3) *Cxcl12* is upregulated in SMCs and ECs in response to increased blood flow; and 4) Analysis of human lung samples from patients with PAH-CHD (VSD) reveals increased expression of CXCL12 in the SMC layer of distal arterioles. This new model will allow the integration of LP/Hx in genetically manipulated mice (including gene knock-out or expression and fate mapping) to explore new pathophysiological mechanisms and potential disease-modifying treatments. Moreover, we provide new insight into the role of Cxcl12 in the development of flow-induced PH.

## Results

### Left pneumonectomy combined with hypoxia results in exacerbated PH and RVH

We hypothesize that exposure to Hx after pulmonary overcirculation (LP) results in exacerbated PH and increased remodeling of distal arterioles (**Fig 1A**). To evaluate this hypothesis, we exposed adult mice to three weeks (wks) of Hx seven days after LP and measured RVSP in experimental mice for comparison (LP/Hx, LP, Hx, and control) (**Fig 1B**). Post-operative day (POD) 14 was chosen as the endpoint for LP/normoxia measurements as our prior experiments demonstrated no significant difference in PH measurements on POD 14 compared to POD 28 and CLG is known to be complete by POD 8^52^. RVSP measurements were elevated in LP/Hx mice compared to Hx (40.4 ±0.9 vs. 33.0 ±0.8mm Hg, *P<*0.0001) and LP mice (27.9 ±0.8mm Hg, *P<*0.0001) (**Fig 1C**). The Fulton index (FI), a measurement of RVH, was also increased in LP/Hx mice compared to Hx (40.0 ±0.8 vs. 31.7 ±0.7%, *P<*0.0001) and LP (31.2 ±0.8%, *P<*0.0001) alone (**Fig 1D**). Hematoxylin and eosin (H&E) staining of the sectioned postmortem hearts in the short axis view revealed a thickened right ventricle (RV) wall after LP/Hx compared to LP and Hx alone (**Fig 1E**). Echocardiogram images of the short axis view in experimental mice also demonstrated RV dilation in end-diastole in LP/Hx mice compared to Hx and LP alone (**Fig 1F**). We have previously shown that RVSP and FI measurements are in a high-normal range at POD 90 after LP^52^. To test the reversibility of our phenotype, we measured RVSP and FI in LP/Hx mice on POD 90 (after they were returned to normoxic conditions) and found similar RVSP measurments (25.2 ±1.8 vs 26.5 ±0.5 mm Hg, *P=*0.59) and FI calculations (26.5 ±1.0 vs 27.0mm Hg ±0.9%, *P=*0.95) compared to our previously published results, demonstrating some long term reversibility of this phenotype (**Fig 1G+H**)^52^.

**Figure 1:**
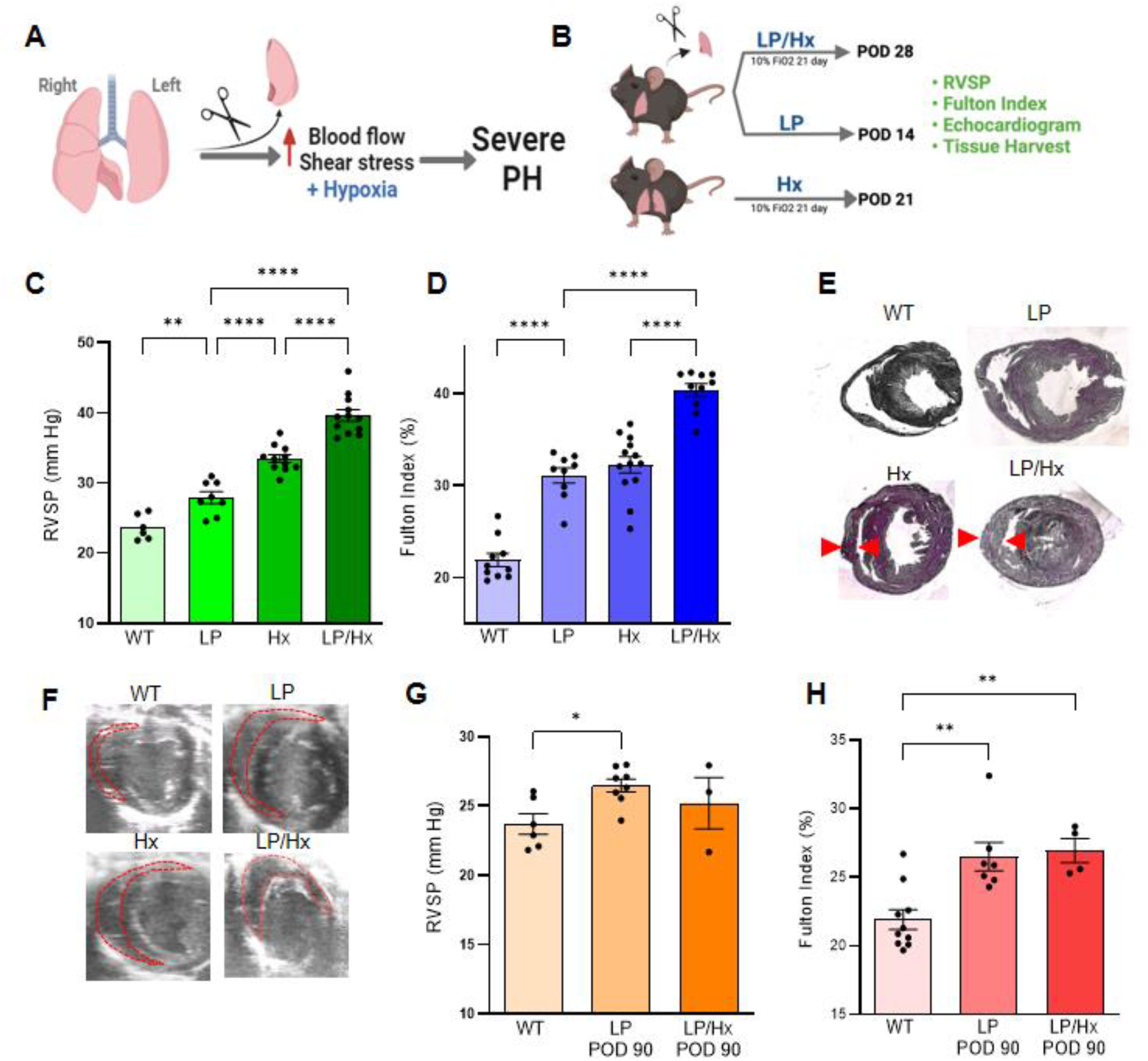
Left pneumonectomy and hypoxia results in exacerbated pulmonary hypertension and right ventricle dysfunction in mice. (A) Illustration showing hypothesis of increased blood flow and Hx resulting in severe PH. (B) Schematic showing experimental design. (C) Right ventricular systolic pressure (RVSP) measurements for control (n=6), LP (n=11), Hx (n=8), and LP/Hx (n=12) mice (D) Fulton index (FI) calculations for RVH in control (n=10), LP (n=13), Hx (n=9), and LP/Hx (n=10) mice. (E) Cross-section of hearts stained with hematoxylin and eosin (H+E) demonstrating thickened RV wall (red arrows) in LP/Hx mice. (F) Representative echocardiogram images of RV in end-diastole with RV outlined in red. (G) RVSP measurements in control (n=6), LP POD 90 (n=11), and recovery mice (n=3). (H) FI measurements in control (n=10), LP POD 90 (n=13), and recovery mice (LP/Hx POD 90) (n=4). Each dot represents an individual mouse sample. Error bars demonstrate mean ± standard error. \**P<*0.05, \*\**P*<0.01; \*\*\**P*<0.001, *P*<*0*.*0001* indicate statistical significance.

### LP/Hx results in exacerbated vascular remodeling and accumulation of SMA on distal arterioles

With the measured hemodynamics revealing LP/Hx mice developed exacerbated PH and RV dysfunction compared to LP or Hx alone, we performed immunofluorescence (IF) staining to quantify the extent of vascular remodeling in the distal arterioles. Whole lung lobe deep tissue clearing and IF staining using iDISCO and light-sheet microscopy demonstrated a significant increase in SMA in the microvessels of LP/Hx mice compared to LP or Hx alone (**Fig 2A**). Next, precision-cut lung slices (PCLSs) were stained for SMA. Confocal microscopy revealed minimal SMA+ cells on the distal arterioles (<50µm) of controls. Notably, all three experimental models (LP, Hx, LP/Hx) revealed an increased accumulation of SMA in the distal arterioles. (**Fig 2B**). Quantification of the distal vasculature from PCLS of LP/Hx mice revealed an increased proportion of small arterioles (<20µm) expressing SMA compared to LP (79.2 ±3.5 vs. 22 ±4.4%, *P<*0.0001) but less so with Hx (72.5 ±0.5%, P=0.51). A similar trend was seen in medium-sized vessels (20-50µm) comparing LP/Hx to LP (80 ±4.4 vs. 47 ±.3.3%, *P<*0.0001) or Hx (70.1 ±1.3%, P=0.21). As expected, there was no difference in SMA amount in large vessels (>50µm) across all four experimental groups (**Fig 2C**). Additional IF staining of PCLSs for ECs (CD31), pericytes (PDGFRb), and SMCs (SMA) was then performed to investigate the contribution of vascular and mural cells to flow-induced vascular remodeling. LP/Hx mice revealed an increased number of cells accumulated on distal arterioles co-expressing PDGFRb and SMA compared to Hx, LP, and control mice (**Figure 2D, top row**). Representative images with increased magnification can be outlined in the yellow boxes (**Figure 2D, bottom rows**).

**Figure 2:**
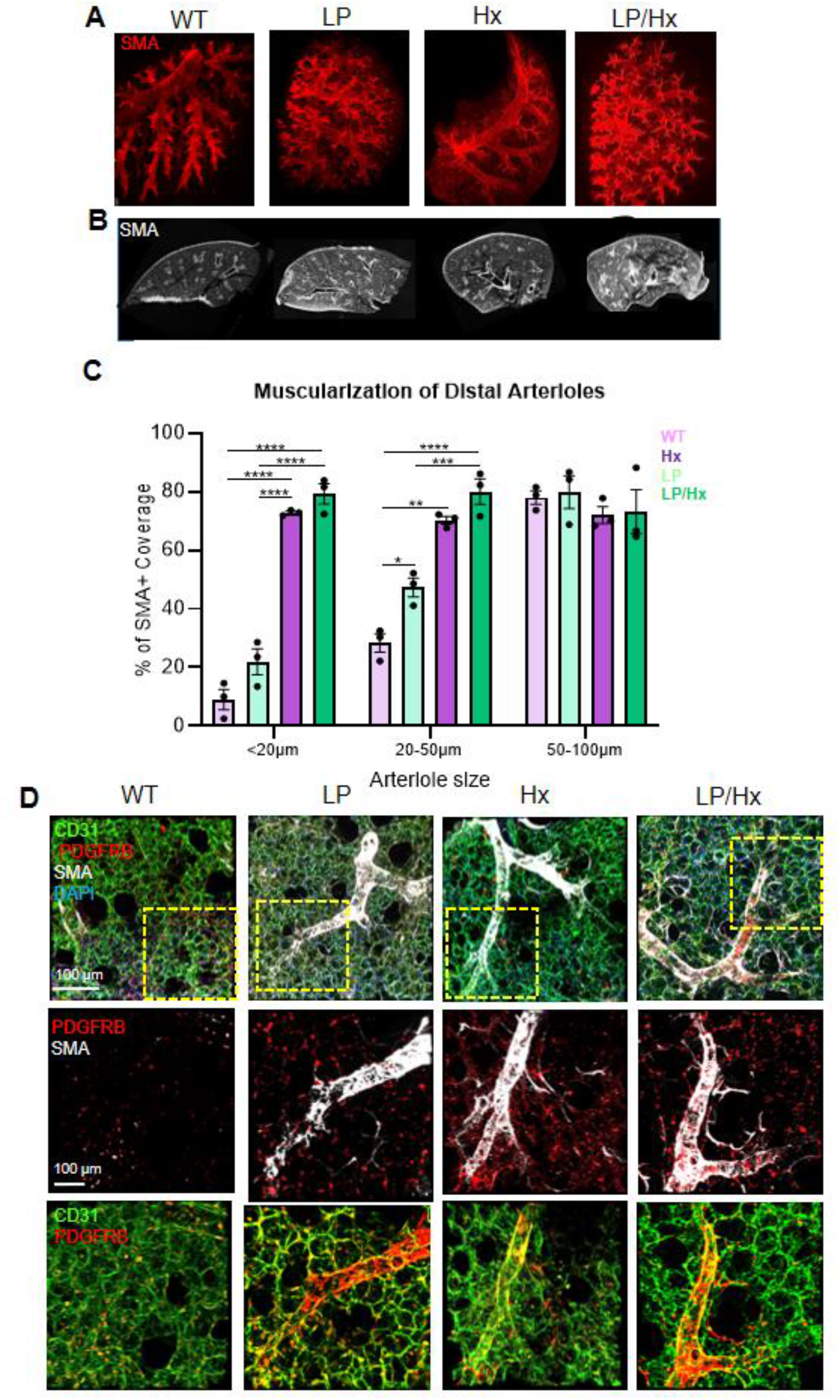
Increased pulmonary blood flow and hypoxia results in exacerbated vascular remodeling and accumulation of SMA on distal arterioles. (A) Whole lung lobe staining of experimental mice for SMA (red). (B) Precision cut lung slices (PCLS) from control, LP, Hx, and LP/Hx mice showing increased accumulation of SMA (white) on arterioles of LP/Hx mice. (C) Graph showing the percentage of distal arterioles (small: <20µm, medium: 20-50µm, large: >50µm) with >50% SMA coverage. Each dot represents one PCLS inspected from a unique animal. N=3 for each experimental group. (D) PCLSs from experimental animals stained for CD31 (endothelium, green), SMA (SMCs, white), PDGFRb (pericytes, red) and DAPI (blue). Scale bar: 100µm. Yellow boxes demonstrate increased magnification. Each dot represents an individual lung sample from a unique mouse. Error bars demonstrate mean ± standard error. *P<0.05; **P<0.01; ***P<0.001; ****P<0.0001 indicate statistical significance.

### Fate mapping using Acta2-CreER::R26-mTmG mice reveals SMC origin cells contribute to increased muscularization on distal arterioles

Due to the increased accumulation of SMA+, PDGFRb+ cells on the distal arterioles of PCLSs from LP/Hx mice, we performed fate-mapping procedures on transgenic mice to further elucidate the roles of SMCs, ECs, and pericytes in vascular remodeling secondary to pulmonary overcirculation. To test our hypothesis and track the response of pulmonary SMCs to increased blood flow and Hx, we utilized *Acta2-CreER::R26-mTmG (*SMA*-mTmG)* mice^54^. After intraperitoneal (IP) injections with tamoxifen, SMCs were appropriately labeled with a green fluorescent protein (GFP). PCLSs were stained for SMA (red) to compare the labeling efficiency. In *SMA-mTmG* control animals, GFP+, SMA+ cells were found almost exclusively in arteries >100µm in size. Responding to LP, Hx, and LP/Hx, there was an increased number of GFP+, SMA+ cells in small arterioles (50µm), indicating SMCs are a source of muscularization and remodeling in flow- and Hx-induced PH (**Fig 3A**). To determine if ECs and pericytes directly contribute to flow-induced vascular remodeling, we performed LP in EC (*Cdh5-CreER::R26-mTmG*, or *Cdh5-mTmG*) and pericyte fate mapping mice (*Cspg4-CreER™::R26-tdTomato*, or *NG2-tdT*)^55-57^. Inspection of PCLSs did not reveal coexpression of ECs (green) or pericytes (red) with SMA staining (**Fig 3B**). Additionally, to confirm the specificity of GFP labeling in SMCs, we performed additional IF staining for fibroblasts with Collagen (COL1A1), Fibronectin (FN), and ECs (CD31) in control and LP/Hx tissue, which demonstrated minimal co-expression of any staining with GFP+ SMCs (**Supplemental Figure 1**). After confirming with fate mapping experiments that SMCs contribute to vascular remodeling in flow-induced PH, we sought to determine the underlying mechanism responsible for these cellular changes.

**Figure 3:**
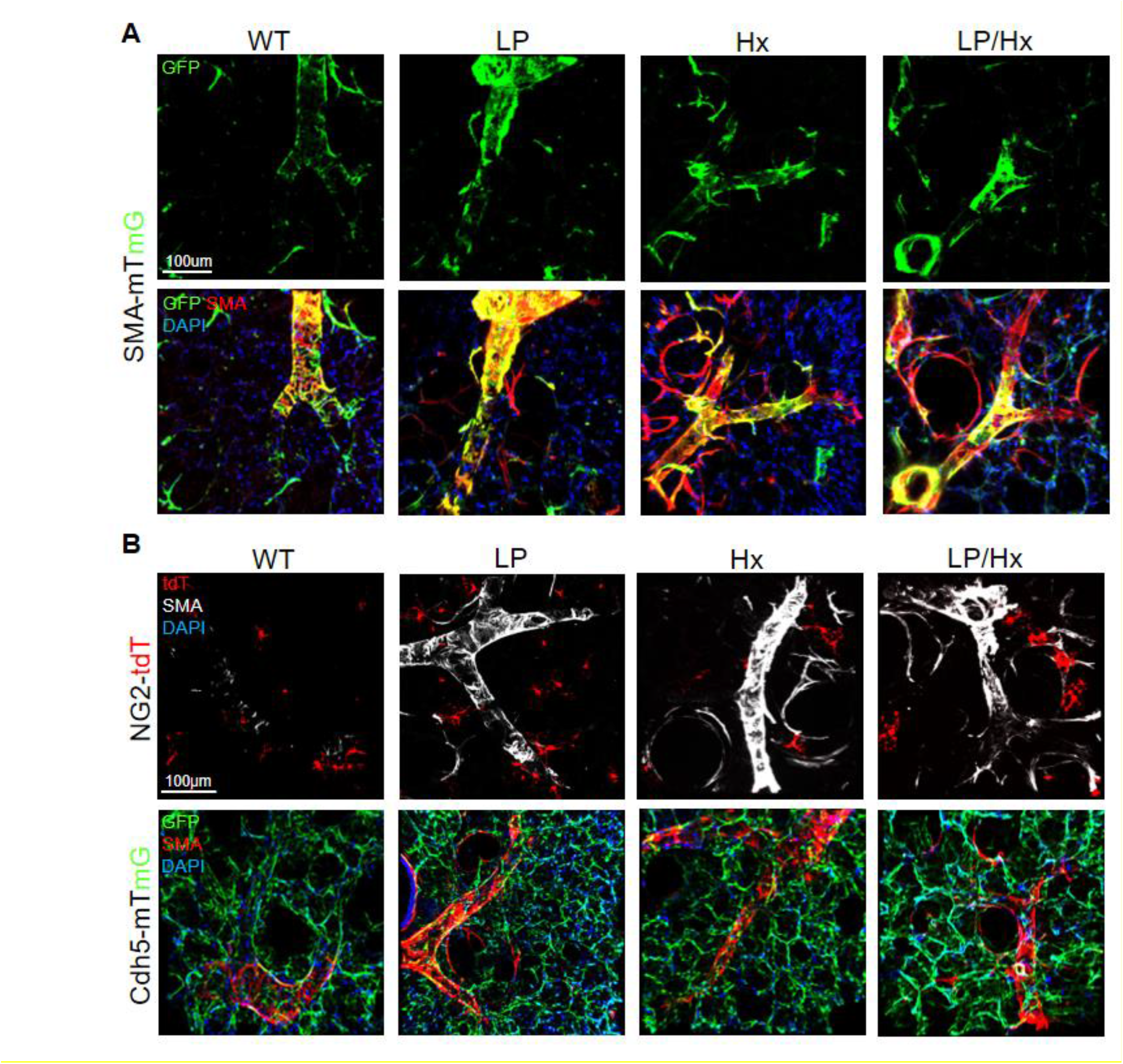
Fate mapping experiments in *Acta2-CreER::R26-*mTmG and *Cdh5-CreER::R26-*mTmG mice reveal abnormal accumulation of SMA on distal arterioles is from SMC origin. (A) Fate mapping experiments of *Acta2-CreER::R26-mTmG* mice in control, LP, Hx and LP/Hx mice. IF staining performed for SMA (red) and DAPI (blue). Notice colocalization of GFP+ cells and SMA staining in LP and LP/Hx mice. Scale bar: 100µm (B) Fate mapping experiments with of *Acta2-CreER::R26-mTmG* mice, *Cdh5-CreER::R26-mTmG* mice and *NG2-CreER::LSL-tdt* mice in control, LP, Hx and LP/Hx mice. IF Staining performed for SMA (red) and DAPI (blue). Scale bar: 100µm

### Pulmonary overcirculation and hypoxia result in the upregulation of Cxcl12 in SMA+ cells and CD31+ cells

After confirming with fate mapping experiments that SMCs contribute to vascular remodeling in flow-induced PH, we sought to determine the underlying mechanism responsible for these cellular changes. There is growing evidence that impaired interactions between ECs and SMCs results in the development and progression of flow-induced vascular remodeling^58,59^. Based on the results from fate-mapping experiments and current knowledge about the role of Cxcl12 in mural cell-mediated vascular remodeling in response to Hx, we isolated SMCs and ECs from mouse lungs using magnetic Dynabeads targeting CD146 and CD31. Real-time PCR (RT-qPCR) from isolated CD146+ SMCs revealed upregulation of *Cxcl12* after LP (1.9-fold increase) compared to controls. SMCs isolated from Hx lungs demonstrated a mild increase in *Cxcl12* compared to LP but were not found to be statistically significant (4.1-vs 1.9-fold increase, *P*=0.15). Interestingly, the LP/Hx model, which developed the most severe PH, RVH, and vascular remodeling compared to LP or Hx alone, demonstrated the highest *Cxcl12* expression compared to control SMCs (6.8-fold increase, LP: *P*=0.007, Hx: *P*=0.08) (**Fig 4A**). CD31+ ECs isolated from three experimental models demonstrated similar results, expression levels of *Cxcl12* were up-regulated 6.5-fold in LP/Hx (LP: *P*=0.0007, Hx: *P*=0.04) while LP had 1.5-fold increase and Hx had a 4.3-fold increase (**Fig 4B**). IF staining of PCLSs from experimental animals revealed that Cxcl12 (white) staining increased around distal arterioles (<50µm) stained with GFP+ SMCs and ECs (**Fig 4C**). Of note, all three experimental mice (LP, Hx, and LP/Hx) demonstrated increased Cxcl12 staining within the lung parenchyma as well, suggesting cell types other than ECs and SMCs may also contribute to Cxcl12 upregulation in response to increased pulmonary blood flow and Hx.

**Figure 4:**
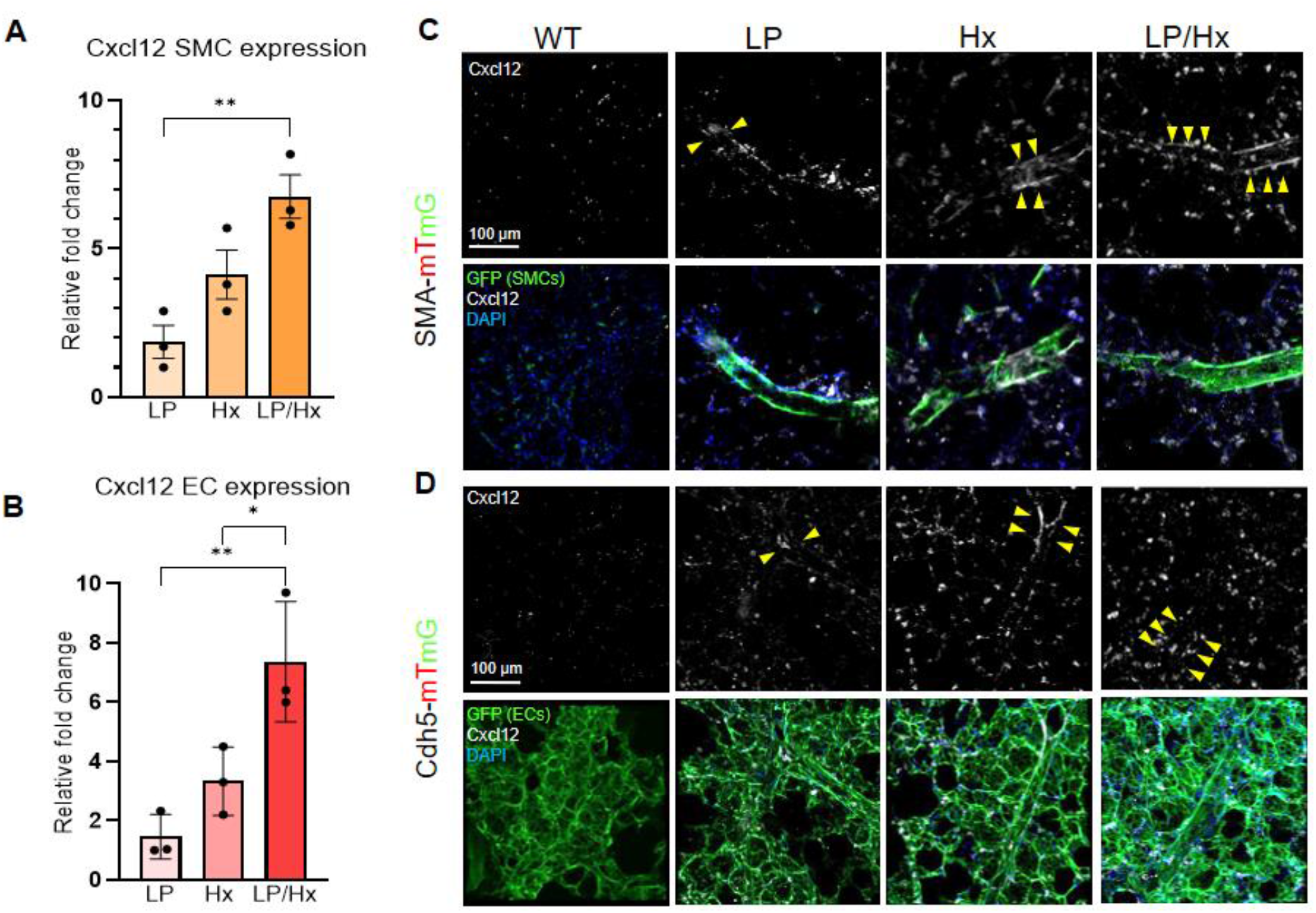
Pulmonary overcirculation and hypoxia results in upregulation of *Cxcl12* in Acta2± and Cdh5± cells. (A) Real-time PCR of isolated SMCs (CD146+) from control, LP, Hx and LP/Hx mice for *Cxcl12*. N=3 for each experimental group. (B) Real-time PCR of isolated ECs (CD31+) from WT, LP, Hx and LP/Hx mice for *Cxcl23*. N=3 for each experimental group. (C) PCLSs from *Acta2-CreER::R26-mTmG* mice showing increased accumulation of Cxcl12 (white) in *Acta2* positive cells (SMCs) and *Cdh5* positive cells (ECs) on distal arterioles (yellow arrows) from lineage tracing mice in control, LP, Hx, and LP/Hx mice. Scale bar: 100µm. Each dot represents a unique sample. Error bars demonstrate mean ± standard error. \**P<*0.05, \*\**P*<0.01; \*\*\**P*<0.001, *P*<*0*.*0001* indicate statistical significance.

### CXCL12 is upregulated in SMCs and ECs from patients with PAH-CHD

To determine if our findings in the LP/Hx murine PH model correlate with human PAH, we performed staining on explant lung samples from non-diseased patients and those with PAH-CHD (PAH secondary to a VSD). Human staining of patients with PAH-CHD was concordant with LP and LP/Hx mice tissue staining, revealing an upregulation of CXCL12 in both SMCs and ECs located within the remodeled distal arterioles compared to non-diseased specimens (**Fig 5A**). Taken together, we provide evidence that pulmonary overcirculation resulted in the upregulation of CXCL12 in SMCs and ECs, which contribute to the vascular remodeling and inflammation seen in human PAH.

**Figure 5:**
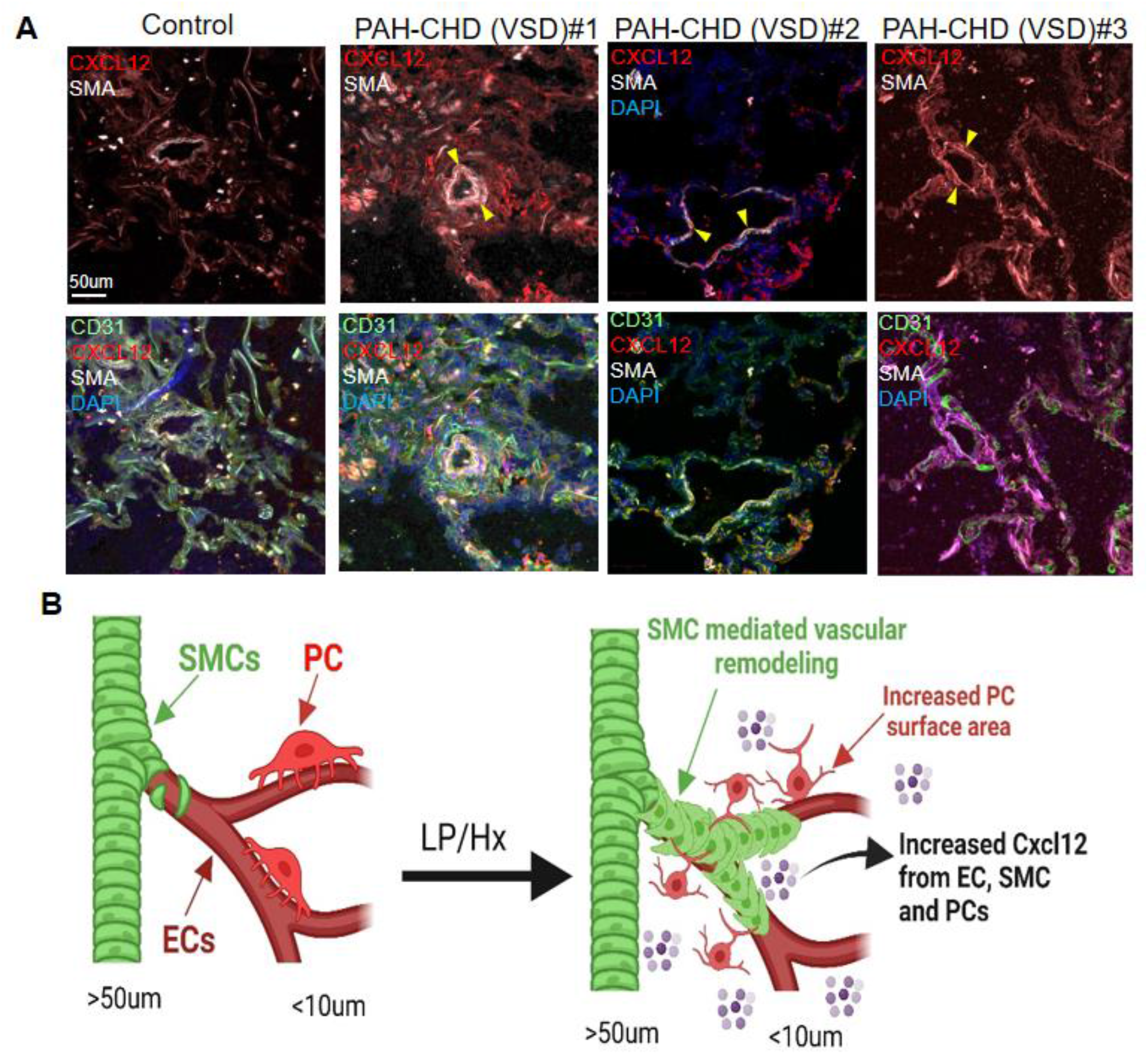
CXCL12 is upregulated in the SMC and EC layer of remodeled vessels in patients with PAH-CHD. (A) Staining of lung tissue from a healthy patient (n=1) and patients with PAH-CHD (VSD) (n=3) for CXCL12 (red), SMA (white), CD31 (green), and DAPI (blue). Yellow arrows highlight CXCL12 accumulation in remodeled distal arterioles. Scale bar: 100µm (B) Schematic illustration showing the role of Cxcl12 and the response of SMCs, ECs, and PCs to increased blood flow and Hx.

## Discussion

Utilizing a combination of linage tracing, IF staining, advanced microscopy, and RT-qPCR, we demonstrated that increased blood flow combined with Hx resulted in exacerbated PH, RVH, and SMC-driven remodeling of distal arterioles with attendant upregulation of Cxl12 (**Fig 5B**). This new murine model of severe PH may better represent the pathophysiological conditions that lead to the vascular remodeling seen in some CHD-PAH patients.

The LP/SUGEN rat model induces severe PH and develops histological features resembling neointimal lesions seen in human disease, while the addition of MCTP in mice results in excessive remodeling and endothelial-to-mesenchymal-cell transition (EndoMT)^21,60 20^. Despite the knowledge gained from these models, the combination of pulmonary overcirculation and EC antagonists does not represent the pathological conditions or inflammatory response that patients go through. Exposure of mice to chronic Hx also results in mild PH and vascular remodeling with increased SMA+ cells accumulated on distal arterioles^25,26^. The contribution of SMCs to vascular remodeling secondary to pulmonary overcirculation is currently unknown, as past studies have focused on the role of ECs and have largely overlooked the contribution of mural cells^20,61^. Herein, we described the cellular changes responding to stimuli (LP and Hx), separately and combined, using three different mouse models (*Acta2-CreER::R26-mTmG, Cdh5-CreER::R26-mTmG*, and *Cspg4-CreER™::R26-tdT*). Results from these experiments demonstrated that SMCs directly contributed to flow-induced vascular remodeling in normoxic and Hx conditions. The molecular mechanism by which SMC-mediated remodeling occurs, whether it is secondary to EC dysregulation, inflammation, or Hx vasoconstriction, is unclear and warrants further investigation. Future studies using the LP/Hx mouse model to describe the communication between mural and other vascular cells will provide novel insights into how these stimuli result in vascular remodeling and the development of PH. Additionally, advanced bioinformatics such as single-cell RNA sequence (scRNA-seq) and assay for transposase-accessible chromatin with sequencing (ATAC-seq) are needed to identify potential pathways that can be targeted for future therapeutic interventions.

Patients with CHD, such as a large VSD, are at risk for the development of PAH. The risk for developing PAH is directly proportional to the size of the defect, and patients with unrepaired VSDs >1.5cm are at a greater risk^62^. Patients with CHD can be exposed to prolonged periods of Hx through associated cardiopulmonary anomalies, sleep-disordered breathing (SDB), infections, and after surgical procedures^30-32,63,64^. Although the exact prevalence of SDB in patients with CHD is unknown, it is suggested to have a higher incidence in patients with CHD and is associated with increased morbidity and mortality^33-35,65,66^. The development of irreversible PH in patients with preexisting pulmonary overcirculation, as seen in certain types of CHD, may result from the additive effect of Hx and increased blood flow, leading to SMC-mediated vasoconstriction and inflammatory/proliferative changes. The LP/Hx murine model of PH can be used in the future to study this hypothesis and the response of cells to two distinct stimuli known to contribute to vascular remodeling: pulmonary overcirculation and Hx. Comparing the molecular pathways in these three models of PH (LP, Hx, LP/Hx) will shed light on the progression and development of PAH seen in patients with CHD and pulmonary overcirculation. Furthermore, it can be utilized as a preclinical model to test the ability of drugs to prevent or treat PH. While studies in rats investigating flow-induced vascular remodeling have histological changes that better represent the lesions seen in patients with PAH compared to mice, they lack the ability to integrate gene manipulation tools such as Cre-LoxP currently only available in mice^25^. Combining the LP/Hx mouse model with gene modification technology will lead to a better understanding of disease pathophysiology and the development of disease-modifying therapies, not previously available in the rat model.

The Cxcl12/Cxcr4 pathway plays a pivotal role in SMC- and pericyte-mediated vascular remodeling in Hx-induced mouse models of PH^39,41,67,68^. Inhibition of the downstream signaling pathway through its receptor, Cxcr4, also leads to decreased SMC proliferation and coverage on distal arterioles^38,39^. The role of Cxcl12 in flow-induced vascular remodeling is unclear. Using IF staining and RT-qPCR, we demonstrated for the first time that isolated SMCs and ECs upregulated *Cxcl12* in flow- and flow/Hx-induced PH. Using SMCs isolated from three different severity of PH models (LP, Hx, LP/Hx), we demonstrated upregulation of *Cxcl12* in SMCs was proportional to the severity of PH. IF staining of biospecimens (explant lung tissues), demonstrated that patients with flow-induced PH (VSD) have upregulation of CXCL12 in the SMC layer of distal arterioles. Identification of the downstream molecules in the Cxcl12/Cxcr4 pathway and relevant inhibitors will lead to future disease-modifying therapies for patients with PAH-CHD. In addition to studying flow-induced PH, LP in mice is a well-established model for studying alveologenesis and CLG. Although the molecular mechanisms and cellular response responsible for stimulating CLG are still unclear, prior studies have shown Cxcl12 is upregulated in the lung parenchyma and drives alveolar regeneration three days after LP ^69^. It is unclear if Cxcl12 promotes abnormal vascular remodeling of distal arterioles, priming ECs for CLG, or both. We demonstrated that *Cxcl12* expression was increased in ECs on POD 14, a time point at which CLG is complete^70-72^. While increased expression of Cxcl12 immediately after pneumonectomy may result in alveologenesis and angiogenesis, increased expression at POD14 suggests its potential inflammatory role in promoting vascular remodeling. It is important to note our study selected specific experimental endpoints commonly used for each rodent model of PH to provide a comparison of hemodynamics. Future studies evaluating mice at alternative and later dates can shed light on the long-term effects of pulmonary overcirculation on vascular remodeling and the reversibility/persistence of proposed PH mouse models.

In conclusion, we established a new and reproducible murine model of severe, flow-induced PH by subjecting mice to 3 wks of Hx after LP. Through lineage tracing and RT-qPCR on isolated SMCs and ECs, we demonstrated that vascular remodeling secondary to increased pulmonary blood flow is mediated by SMCs and associated with the upregulation of Cxcl12. By integrating this model with gene knock-out and additional fate mapping strains, along with advanced bioinformatics approaches, researchers can investigate the molecular mechanisms and cellular responses underlying flow-induced PH.

## Figures

**Supplementary Figure 1:**
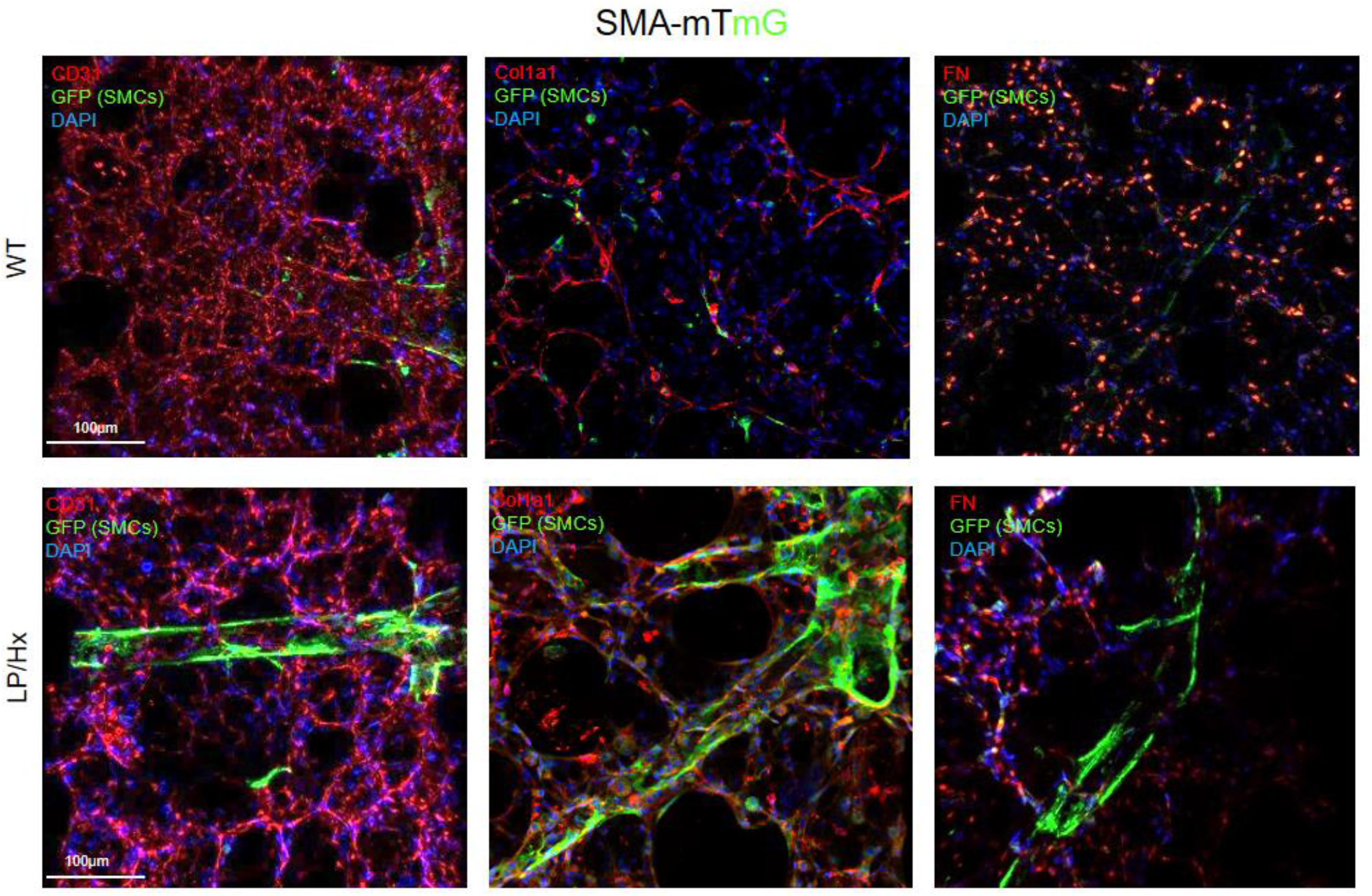
Control and LP/Hx *Acta2-CreER::R26-*mTmG mice do not demonstrate Acta2+ cells with fibroblast and EC markers. PCLSs and IF staining in *Acta2-CreER::R26-mTmG* control and LP/Hx mice for ECs (CD31, red) collagen (COL1A1, red), fibronectin (FN, red), and DAPI (blue). Scale bar: 100µm.

## Materials

### Experimental Procedures

Studies in this paper were approved by the Animal Care Committee and Institutional Guidelines of Boston Children’s Hospital. All data is contained within the manuscript and raw images or additional information can be shared upon request from the corresponding author.

### Animals

Experiments adhered to the National Institutes of Health guidelines use of laboratory animals. *Acta2-CreER::R26-mTmG* mice (C57BL/6J) were used for SMC fate mapping experiments, *Cdh5-CreER::R26-mTmG* mice for EC fate mapping, and *Cspg4-CreER™-tdTomato* for pericyte fate mapping. Ear clip-based genotyping was used to identify experimental mice.

Male mice were six wks of age or older at the time of tamoxifen surgery or exposure to Hx. Comparison of hemodynamic data of LP, LP at POD 90, and control mice was utilized from our group’s previous publication^52^.

### Tamoxifen administration

*Acta2-CreER::R26-mTmG* and *Cdh5-CreER::R26-mTmG* mice were administered 10mg of tamoxifen via intraperitoneal (IP) injection (20 mg/ml dissolved in corn oil) over five days *Cspg4-CreER™-tdTomato* mice were injected with 2mg of tamoxifen once. All mice were at least 5 weeks of age before injection. Transgenic mice were allowed to rest for a minimum of seven days before undergoing pneumonectomy or being exposed to Hx.

### Hypoxia studies

For chronic Hx experiments, mice were exposed to 10% FiO_2_ via Hx chamber with ad lib access to food and water for 3 wks. The chamber was maintained with a continuous mixture of room air and nitrogen gas. The chamber environment was monitored using an oxygen analyzer (Servomex, Sugar Land, TX) and CO_2_ was removed with lime granules. The internal chamber was monitored daily for O_2_ concentration, CO_2_ concentration, humidity, and animal welfare. To establish the LP/Hx mouse model, mice were exposed to 3 wks of Hx seven days after LP. RVSP measurements and tissue harvest were performed after exposure to Hx, or on POD 28.

### Left pneumonectomy

Animals were anesthetized via IP injection with a mixture of ketamine (80-100 mg/kg) and xylazine (5-10 mg/kg). After reaching an appropriate level of sedation, mice were placed on a vertical platform and a light-source was used to transilluminate the vocal cords. Intubation was achieved with a 22-gauge angiocatheter (Becton Dickinson, Franklin Lakes, NJ) and animals were ventilated at 180 breaths/min using a rodent ventilator (MiniVent Ventilator; Harvard Apparatus, Holliston, MA). After adequate hair removal to the animals left thorax and appropriate asepsis technique, a left anterior mid-axillary incision was made extending from the axilla to the costal margin^74,75^. The thoracic cavity was entered at the 5^th^ intercostal space and the left lung was carefully lifted through the incision and ligated using a 4-0 silk suture (Ethicon, Raritan, NJ). The rib cage and thoracic cavity were then closed with a 4-0 PDS suture (Ethicon, Raritan, NJ). The skin was closed using a 4-0 nylon suture (Ethicon, Raritan, NJ). Mice were provided with appropriate analgesia and monitored daily for signs of pain and respiratory distress during the post-operative period.

### Mouse lung tissue clearing and preparation of precision cut lung slices

After euthanasia, mice were placed in the supine position and the abdominal and thoracic cavity exposed through a midline incision. The sternum was then dissected to expose the contents of the mediastinum. The abdominal aorta was then located and cut. Next, a butterfly needle was inserted into the right ventricle, and the heart and lung slowly perfused with 15cc of 1x PBS to flush the red blood cells from the circulatory system. Once the lung lobes were white in appearance, the trachea was located and cannulated. Lungs were inflated with 2% low-melting point agarose. After adequate inflation, the trachea was tied and the mediastinal contents removed and placed in ice-cold 1x PBS to solidify the agarose. The lung tissue was then fixed in 4% paraformaldehyde (PFA) at 4°C overnight, followed by washing in 1x PBS for an additional day. Lung lobes were then separated and sectioned (Leica VT1000 S) at a thickness of ∼300µm for future IF staining experiments and confocal microscopy.

### Morphometric analysis of vascular remodeling

PCLS of experimental mice (control, LP, Hx, LP/Hx) underwent IF staining for endothelium (CD31) and smooth muscle actin (SMA). One stained tissue from each mouse was scanned with a confocal microscope (880 laser scanning confocal with Fast Airyscan) using the Z-stack application to visualize the three-dimensional structure and anatomy of pulmonary vasculature (CD31+ staining) of various diameters (<20µm, 20-50µm and >50µm). Once identified, the vessel was inspected to determine the degree of SMA coverage (greater than 50% of vessel co-stained with SMA). PCLS were scanned from top to bottom with all areas of tissue inspected. The percentage of arterioles expressing SMA for each group was reported for all experimental conditions (N=3 per group).

### Immunofluorescence Staining

PCLSs were blocked with 5% goat or donkey serum in 0.5% Triton X-100/PBS (PBS-T) for one hour. Samples were then incubated in the dark with primary antibodies for one to two days at 4°C. After incubation, tissue was washed three times in 1x PBS followed by incubation with secondary antibodies overnight in the dark at 4°C. PCLSs were again washed and placed on microscope slides with mounting media containing DAPI (Vector Labs). All images were captured using a Zeiss confocal 880 Airyscan 2 and processed by Aivia software. The following antibodies were used for human and mouse lung tissues:

Mouse-anti-mouse/human SMA-647 (1:50; Santa Cruz, sc-32251)

Rat-anti-mouse CD31(1:100; BD-Pharmingen, 553370)

Rabbit-anti-mouse PDGFRb (1:200; Invitrogen, MA5-15143)

Rabbit-anti-mouse/human SDF-1 (1:200; Abcam, ab9797)

Sheep-anti-human PECAL-1 (1:100. R&D Systems, BAF806)

Rabbit-anti-mouse/human COL1A1 (1:200; Cell Signaling Technology, 39952)

Rabbit-anti-mouse Fibronectin-647 (1:100; Cell Signaling Technology, 26836)

### Heart OCT Compound

Hearts were removed from the mediastinum after sacrifice and washed thoroughly in 1x PBS. Heart samples were then transferred to 4% paraformaldehyde (PFA) overnight at 4°C. The next morning, samples were washed in 1x PBS for 24 hr and placed in 30% sucrose for a minimum of 24 hr. After sitting in sucrose, samples were embedded in OCT gel and flash-frozen using dry ice. The frozen samples were stored at -80°C until the were sectioned with a cryostat machine into thin slices. Samples were sectioned until RV and left ventricle opening became visible and then mounted on microscopy slides for hematoxylin and eosin staining. Images were captured with a light microscope (Olympus BX-41 upright microscope, Olympus) at 4× objective.

### Whole mount lung lobe clearing and immunofluorescence staining

After removing the lung from appropriately euthanized animals, the lung lobes were fixed with 4% PFA, washed in PBS containing 0.2% Trition X-100 (PTx.2) and then incubated in 1x PBS/0.2% TritionX-100/20% DMSO overnight. Samples were submerged in 1xPBS/0.1%Tween-20/0.1%Tritonx-100/0.1%Deoxychlorate/0.1%NP40/20% DMSO overnight. The next day, samples were washed with PTx. 2 and placed in permeabilization solution (PTx.2/2.3% glycine/20% DMSO) for 48 hours. Samples were incubated in blocking solution (PTx.2/10% DMSO/5% normal goat serum) for two days and then submerged in primary antibodies diluted in PTx.2/0.001% heparin (PTwH)/5% goat serum/10% DMSO for a minimum of three days. After incubation, samples were washed in PTwH and placed in secondary antibodies diluted in PTwH/5% goat serum for three days. After secondary staining, lung lobes were washed in PTwH and progressively dehydrated in 100% methanol over two days and then placed in 33% methanol/66% dichloromethane for three hours. Finally, lung samples were washed in dichloromethane and then cleared in dibenzylether (DBE). Images were obtained using a light sheet microscope.

### Right ventricle systolic pressure and Fulton index measurements

To calculate right ventricle systolic pressure (RVSP) measurements, mice were anesthetized using isoflurane (2.0% in 2LPM O_2_) administered through a nose cone. After reaching an appropriate level of anesthesia, a small incision was made in the skin to the right of the trachea. The surrounding fat was removed with blunt dissection to isolate and visualize the right jugular vein. The vein was then cannulated using a 1.4F catheter (Millar Instruments) and carefully and slowly advanced into the right ventricle. The RVSP tracings were analyzed using PowerLab software. RVSP measurements for each animal was obtained by calculating measurements at three different time points separated by a minimum of 30 seconds.

Once RVSP measurements were obtained, mice were euthensized with cervical dislocation. The entire heart was removed from the chest cavity and the atrium and pericardial fat dissected away. The interventricular septum between the right and left ventricle was then identified, and the two ventricles carefully separated. The weight of the right ventricle and left ventricle with the interventricular septum was obtained. The Fulton index was calculated using the following equation: (Right Ventricle Mass)/(Left Ventricle Mass + Septum Mass).

### Echocardiography

Images of the right ventricle in experimental animals were obtained using transthoracic echocardiogram with VisualSonics Vevo 3100 Software (VisualSonics Inc. Toronto, ON, Canada) and a small rodent transducer (MS-550D, 22–55MHz). To obtain images, mice were first anesthetized in an induction chamber filled with 2-3.0% isoflurane and 100% oxygen. Appropriately anesthetized mice were then placed in the supine position and a nosecone used to deliver 1.5-2.5% isoflurane mixed with oxygen (1.0 L/min) on a heated stage while echocardiogram was performed. The chest fur of animals was completely removed with a chemical hair remover. Representative echocardiogram images of the RV were obtained in the short-axis view in end diastole

### Cell isolation

Lung tissue was harvested from three experimental mice of the similar condition, combined, and digested using commercial Miltenyi gentle MACS dissociator and lung tissue dissociation kit (Miltenyi Biotec, Germany). After digestion, 10 mL of 0.1% FBS/PBS was added to the cell suspension and the solution was passed through a BD Falcon 70μm strainer (BD Biosciences, San Jose, CA) to remove any undigested tissue. The solution was centrifuged at 400G for five minutes and the cell pellet then resuspended in 1 mL 1x PBS containing CD45 IgG– coated magnetic Dynabeads and the cell and beads gently rotated at 4°C for 30 minutes to deplete immune cells. CD45+ dynabeads were then removed by placing the sample on a magnetic stand and the remaining cells (CD45-) were incubated in CD140b-APC antibody (Miltenyi Biotec, Germany) for 30 minutes at 4°C on a shaker. Cells were washed with 1x PBS and then stained with anti-APC beads (Miltenyi Biotec, Germany) and allowed to incubate for another 30 minutes at 4°C on a shaker. The Miltenyi Biotech Automacs Cell Sorter was then used to isolate CD140b+ cells, which were then digested in 500 µL of Trizol for RNA isolation. CD45-, CD140b-cells were then resuspended in 1mL of 1x PBS containing CD146 IgG-coated Dynabeads and gently rotated for 30 minutes at 4°C. CD146+ beads were then selected with magnetic dissociation, washed with 1x PBS and digested in 500uL Trizol. The remaining cell solution (CD45-, CD140b-, CD146-) were resuspended in CD31-IgG coated Dynabeads and again rotated for 30min at 4°C. The CD31-beads were then washed with 1x PBS, selected with magnetic dissociation and digested in 500ul Trizol for RNA isolation.

### Real-time Quantitative PCR

mRNA expression was determined by quantitative Taqman real-time PCR. Cellular RNA was extracted using an RNeasy Kit (Qiagen, Redwood City, CA) and converted to cDNA using the High-Capacity cDNA Reverse Transcription Kit with RNase Inhibitor (Thermo Fisher Scientific, Waltham, MA) following the manufacturer’s protocol. All real-time quantitative PCR studies were run in triplicates. ΔCT determined the difference in mRNA expression against housekeeping genes (GAPDH, B2M, 18S). Primers were purchased from Integrated DNA Technologies.

### Statistical analysis

The number of animals per experiment can be found in the figure legends. All data is expressed as mean ± standard error of mean (SEM) unless otherwise indicated. Statistical significance was determined using either an unpaired t-test or ordinary two-way ANOVA. \**P<*0.05, \*\**P<*0.01, \*\*\**P<*0.001, \*\*\*\**P<*0.0001.

## Acknowledgment/Funding

The content is solely the responsibility of the authors and does not necessarily represent the official views of the National Institutes of Health.This manuscript was supported by the following grants: NINDS P30 Core Center Grant #NS072030 from Harvard NIF core for imaging or analysis; National Institutes of Health 1R01HL150106, ATS/PHA Aldrighetti Research Award for Young Investigators, Bayer PHAB award (KY), 2T32DK007754-22 (STT), 5T32HL007633-36, F32HL167554-01A1, and the ATS Early Career Investigator Award in Pulmonary Vascular Disease (TK), Boston Children’s Vascular Biology Program and The Boston Children’s Hospital Surgical Foundation (MP). The authors acknowledge the BioRender program which was used for the illustrations in this manuscript.

## Author contributions

**Timothy Klouda**: Conceptualization, Methodology, Formal Analysis, Investigation, Writing-Original Draft, Visualization, Project administration. **Savas T. Tsikis**: Conceptualization, Methodology, Investigation, Writing-Review & Editing. **Thomas I. Hirsch**: Conceptualization, Methodology, Investigation, Writing-Review & Editing, **Yunhye Kim**: Investigation, Writing-Review & Editing. **Tiffany Liu**: Investigation, Writing-Review & Editing. **Ingeborg Friehs**: Writing-Review & Editing. **John Y.-J. Shyy**: Writing-Review & Editing. **Gary Visner**: Writing-Methodology, Review & Editing. **Benjamin A Raby**: Conceptualization, Writing-Review & Editing. **Mark Puder**: Resources, Writing-Review & Editing, Supervision, Funding acquisition. **Ke Yuan**: Conceptualization, Methodology, Investigation, Resources, Writing-Original Draft, Supervision, Funding acquisition

## Abbreviations

ATAC-seq: Assay for transposase-accessible chromatin with sequencing
CHD: congenital heart disease
CLG: compensatory lung growth
EC(s): endothelial cells
EndoMT: endothelial to mesenchymal cell transition
GFP: green fluorescent protein
Hx: hypoxia
IF: Immunoflourescence
LP: left pneumonectomy
MCT: monocrotaline
MCTP: monocrotaline pyrrole
RV: right ventricle
RVH: right ventricle hypertrophy
FI: Fulton index
RVSP: right ventricle systolic pressure
RV: right ventricle
PCLS: precision cut lung slice
PAH: pulmonary arterial hypertension
PAP(s): pulmonary artery pressures
PH: pulmonary hypertension
POD: postoperative day
RT-qPCR: real-time qPCR
Single-cell RNA sequence: scRNA-seq
SDB: sleep-disordered breathing
SMC(s): smooth muscle cells
Sugen-5416: SUGEN
VSD: ventricular septal defect
Week: wk(s)

